# Maize *YABBY drooping leaf* genes regulate floret development and floral meristem determinacy

**DOI:** 10.1101/395186

**Authors:** Josh Strable, Erik Vollbrecht

**Affiliations:** Department of Genetics, Development and Cell Biology, Iowa State University, Ames, Iowa 50011; Interdepartmental Plant Biology, Iowa State University, Ames, Iowa 50011

## Abstract

Floret units in cereals produce grain, directly impacting yield. Here we report mutations in the maize *CRABS CLAW* (*CRC*) co-orthologs *drooping leaf1* (*drl1*) and *drl2* alter the development of ear and tassel florets. Pistillate florets of *drl1* ears appear sterile and display ectopic unfused carpels that fail to enclose an expanded nucellus. Staminate florets of *drl1* tassels have extra stamens and retain fertile anthers. Natural variation and transposon alleles of *drl2* enhance *drl1* floret phenotypes by reducing floral meristem (FM) determinacy. The *drl* paralogs are co-expressed in lateral floral organ primordia, but not within the FM. Together, the expression patterns and indeterminate mutant FMs suggest that the *drl* genes regulate FM activity and impose meristem determinacy by a non-cell autonomous signal. Genetic interaction analyses of *drl* mutants with maize floral mutants indicate that the *drl* genes are required throughout floret development, illustrating their importance for proper floret patterning in maize.

## Introduction

A major goal in plant biology is to understand the factors that regulate meristem activity. Meristems, active, pluripotent stem cell tissues, produce all postembryonic organs of flowering plants^1^. Meristem determinacy (degree of meristem activity) is a critical factor that shapes vegetative, inflorescence and floral architectures. Vegetative and inflorescence meristems are indeterminate, producing an unspecified number of lateral primordia. Floral meristems (FMs) are generally determinate, initiating a more set number of floral whorls and organs before undergoing terminal differentiation. Commonly, eudicot flowers are composed of four whorls of floral organs (outermost to innermost: sepal, petal, stamen, and carpel). Similarly in the monocots, grass florets (flowers), including those in maize, are arranged in whorls of floral organs, some of which have grass-specific names (outermost to innermost: lemma, palea, stamen and carpel). As each grain is the product of one floret, regulating FM activity is a key agronomic trait.

FMs pattern flowers through the combinatorial activity of four classes of gene functions that dictate organ identity and FM determinacy^2,3^. In Arabidopsis, carpels are specified by the MADS-box transcription factor AGAMOUS (AG)^4^. Additionally, AG controls FM determinacy by repressing expression of the stem cell regulator *WUSCHEL*^5,6^. In the cereals, where AG orthologs have expanded and undergone subfunctionalization during grass evolution, FM determinacy is regulated redundantly^7-9^. The maize *AG* ortholog *zea agamous1* (*zag1*) imposes FM determinacy with a lesser obvious role in regulating floret organ identity as supernumerary carpels develop in pistillate florets of *zag1* mutants^7^. ZAG1 interacts physically with *AG-LIKE6* (*AGL6*) subfamily member BEARDED EAR (BDE)/ZAG3, and *zag1; bde* double mutants reveal a synergistic interaction in regulating FM determinacy^10^. The maize *indeterminate floral apex1* (*ifa1*) mutant affects FM determinacy in pistillate and staminate florets, and in the innermost staminate whorl of *ifa1* mutants, a nucellus develops at very low penetrance^11^. *ifa1* shows a redundant genetic interaction with *zag1* to regulate FM determinacy^11^. In the bisexual florets of rice, AG orthologs *OsMADS3* and *OsMADS58* regulate floret organ identity and FM determinacy, respectively^8,9^.

Arabidopsis *YABBY* family member *CRC* is required for proper growth of the gynoecium^12^. Loss-of-function mutations in *CRC* consistently result in reduced stylar growth and incomplete medial fusion of carpels. *crc* mutants occasionally produce three carpels compared with two in wild-type, suggesting *CRC* is necessary to promote FM determinacy^12,13^. Expression of *CRC* is restricted to developing carpels and nectaries^13^. The rice *CRC* ortholog *DROOPING LEAF* (*DL*) is required for carpel identity, as carpels undergo homeotic transformation to stamens in loss-of-function *dl* mutants^14^. Transformed stamens are variable in number, indicating *DL* also regulates FM determinacy. *DL* is expressed in carpel primordia of rice florets^14^. Genetic analysis indicates that *DL* and the rice *AGL6* subfamily member *MOSAIC FLORAL ORGAN1* (*MFO1*)/*OsMADS6* redundantly regulate FM determinacy^15^.

Here we report the maize *CRC* co-orthologs *drl1* and *drl2* are required for the development of dimorphic, unisexual ear and tassel florets. *drl1* pistillate florets are sterile, display ectopic unfused carpels and have an expanded nucellus. *drl1* staminate florets have extra stamens with fertile anthers. Natural variation and transposon alleles of *drl2* enhance *drl1* floret phenotypes by further reducing FM determinacy. The *drl* paralogs are co-expressed in lateral organ primordia initiated by the FM, but not within the FM, and interact genetically with other maize floral mutants. Our results demonstrate that the *drl* genes are required for floret patterning and to impose meristem determinacy via some non-cell autonomous mechanism, and thus, provide critical control of FM activity in the development of grain-bearing structures.

## Results

### *drl1* and *drl2* regulate floret development

Maize staminate and pistillate florets are produced on the tassel and ear, respectively^16^. During tassel and ear development, branching events from multiple meristem types^17^ ultimately give rise to floret whorls housed in grass-specific spikelets^18^. The indeterminate inflorescence meristem (IM) of the tassel and ear initiates determinate spikelet pair meristems (SPMs); additionally in the tassel, the IM initiates indeterminate branch meristems (BMs). Each SPM produces a pair of determinate spikelet meristems (SMs), each of which gives rise to two glumes; afterwards, each SM initiates a lower floral meristem (LFM) and then converts identity to an upper floral meristem (UFM). Each determinate LFM and UFM gives rise to a lemma, a palea, two lateral-abaxial lodicules, three stamens and three carpels^19^. Two lateral-adaxial stamens are spaced widely relative to the medial-abaxial stamen^20^. Three connately fused carpels form the single pistil; however, only the two lateral-abaxial carpels (indeterminate carpels, C_i_) form an elongated silk, whereas growth of the medial-adaxial carpel (determinate carpel, C_d_) is limited in growth to envelop the single ovule^21,22^. The ovule consists of nucellus tissue enclosed mostly by inner and partially by outer integuments, all within a locule formed by the three fused carpels^16^. After organ initiation, sex determination in the tassel and ear culminates in abortion of the carpel whorl in staminate florets and arrest of stamen primordia in pistillate florets, respectively (Fig. 1a,b)^19^.

**Figure 1.**
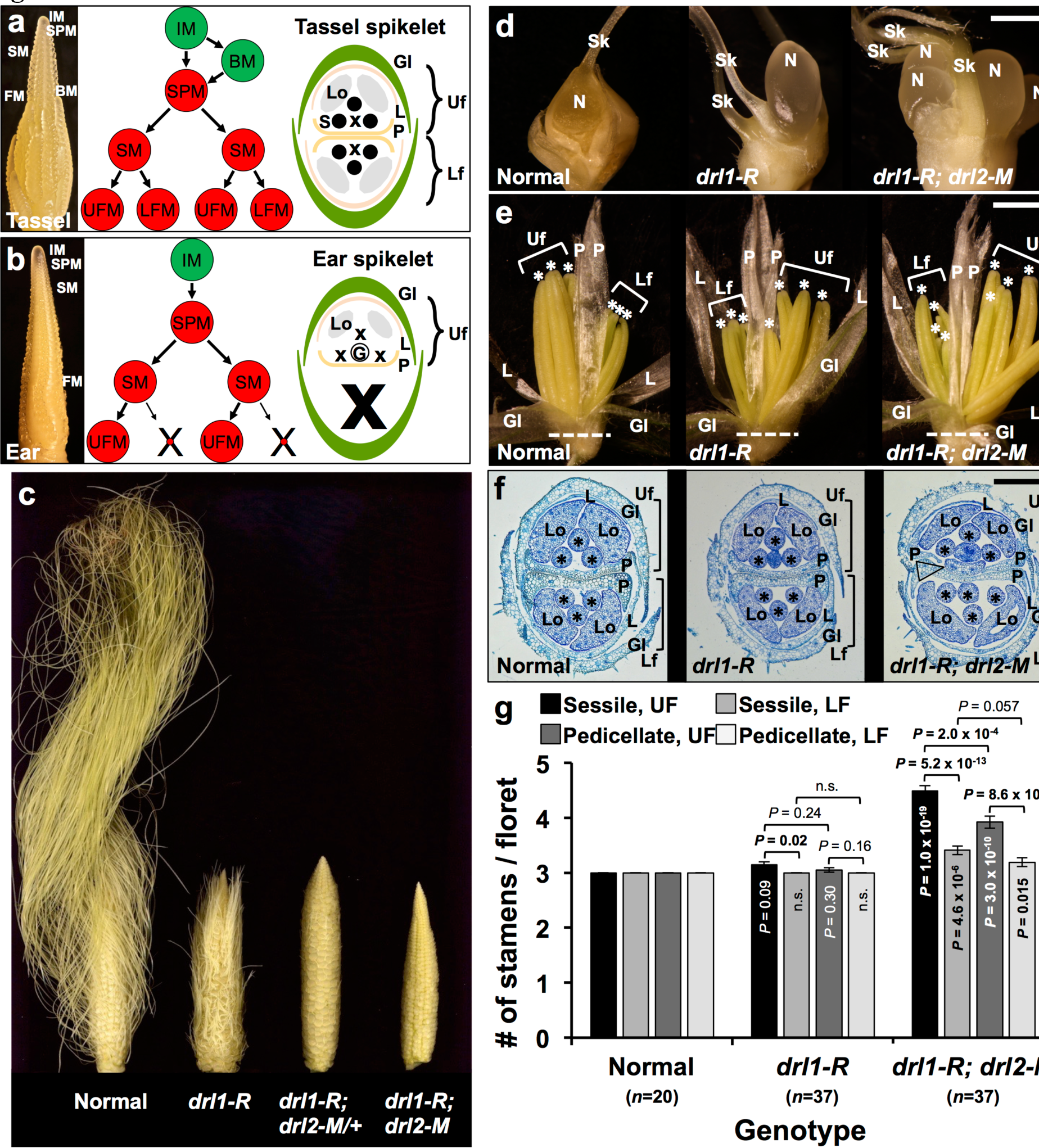
Floral architecture in *drl1* single and *drl1; drl2* double mutants. **a,** Development of the maize tassel and staminate florets. Left, developing tassel (6 mm). Middle, the IM gives rise to BMs and SPMs, which in turn produce SMs that then give rise to UFM and LFM. Right, a tassel spikelet consists of outer and inner Gl that house the UF and LF, each of which is comprised of a Le and P, two Lo, and three S. **b,** Development of the maize ear and pistillate florets. Left, developing ear (6 mm). Middle, the IM gives rise to SPMs that produce SMs, which give rise to UFM and LFM; the LFM aborts early. Right, an ear spikelet consists of outer and inner Gls that house the UF comprised of a Le and P, two reduced Lo, and a G. **c,** Mature ears from an F_2_ population showing dosage effects of the *drl* loci: left to right, normal, *drl1-R*, *drl1-R; drl2-M/+* and *drl1-R; drl2-M*. **d,** Dissected pistillate spikelet of normal (left, bisected), *drl1-R* (middle, intact) and *drl1-R; drl2-M* (right, intact). **e,** Dissected staminate spikelet of normal (left), *drl1-R* (middle) and *drl1-R; drl2-M* (right). **f,** Transverse sections of staminate spikelets stained with Toluidine Blue-O taken at the plane of the dashed line in (**e**); normal (left), *drl1-R* (middle) and *drl1-R; drl2-M* (right). **g,** Quantification of stamens in normal, *drl1-R* and *drl1-R; drl2-M* florets. Mean ±SEM, *P*-values based on two-tailed Student’s *t* tests; *n*, sample size. IM, inflorescence meristem; BM, branch meristem; SPM, spikelet pair meristem; SM, spikelet meristem; UFM, upper floral meristem; LFM, lower floral meristem; Gl, glume; L, lemma; P, palea; Lo, lodicule; S, stamen, G, gynoecium; X, aborted organ; UF, upper floret; LF, lower floret; N, nucellus; Sk, silk. Asterisks mark mature anthers. Arrowhead marks ectopic organ.Scale bars, 2 mm (**d**,**e**) and 200 µm (**f**).

Likely null mutations of *drl1* displayed aberrant pistillate and staminate floret morphologies. Macroscopically, *drl1* mutant ears appeared sterile, with underdeveloped silks consisting of reduced, unfused carpel walls that failed to enclose an expanded nucellus (Fig. 1c,d). *drl1* pistillate phenotypes were reminiscent of the floret phenotypes described for the *ifa1* mutant^11^. We found *drl1* and *ifa1* to be allelic through genetic complementation (Supplementary Fig. 1) and by sequencing the *drl1* locus in *ifa1* mutant plants^23^. Additionally, we observed florets with higher-order branching in *drl1-R; zag1-mum1* double mutants, where floret axes displayed iterative secondary and tertiary branch-like lateral growth from the axils of ectopic palea or bracts (Supplementary Fig. 2), corroborating a previously reported synergistic interaction in *ifa1; zag1-mum1* double mutant florets^11^. *drl1* and its paralogous genetic enhancer locus, *drl2*, encode *CRC* co-orthologs in the *YABBY* family of transcriptional regulators^23^.

**Figure 2.**
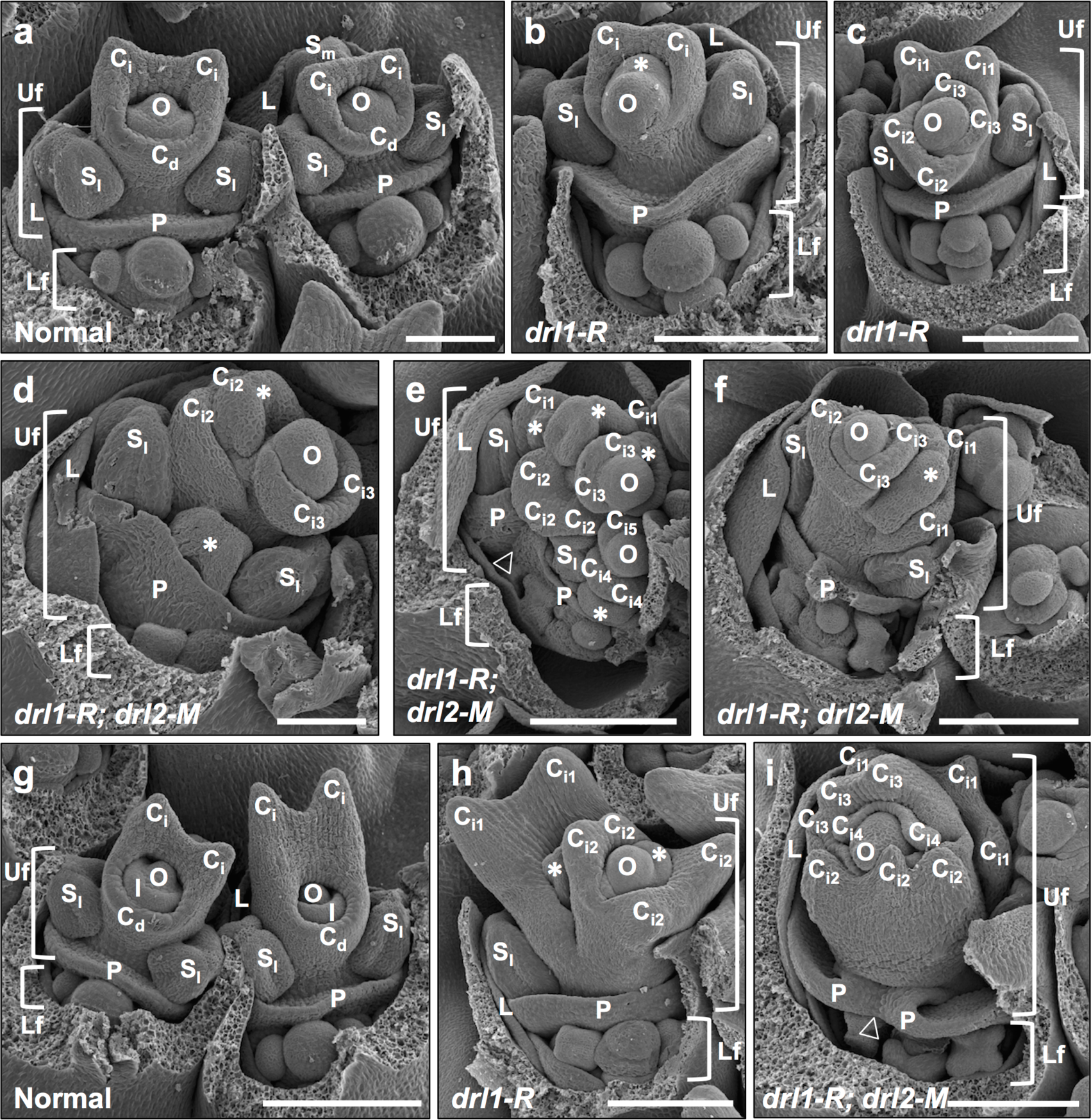
Development of pistillate florets in *drl1* single and *drl1; drl2* double mutants. **a-f,** Scanning electron micrographs (SEMs) of mid-staged pistillate florets from normal (**a**), *drl1-R* (**b,c**) and *drl1-R; drl2-M* (**d-f**) developing ears. **g-i,** SEMs of late-staged pistillate florets from normal (**g**), *drl1-R* (**h**) and *drl1-R; drl2-M* (**i**) developing ears. Glumes were removed manually to expose the upper and lower florets. Uf, upper floret; Lf, lower floret; L, lemma primordium; P, palea primordium; S_l_, lateral stamen primordium, S_m_, medial stamen primordium, C_i_, indeterminate carpel primordium; C_d_, determinate carpel primordium; I, integument; O, ovule primordium. C_i_ whorl number is subscripted. Asterisks (**b,d,e,h**) mark ectopic primordia. Arrowhead (**i**) points to palea involution. Scale bars, 100 µm (**a,d**), 200 µm (**b,c,e-i**).

Genetic combinations between *drl1* alleles and the loss of- or low-function *drl2-Mo17* natural variant allele (hereafter referred to as *drl2-M*) or the strong *drl2-DsD08* transposon allele^23^ enhanced all aspects of the *drl1* floret phenotype, such that florets from double mutants displayed multiple, expanded nucelli that appeared to originate from sustained FM activity in the upper floret (UF) (Fig. 1c,d and Supplementary Fig. 3). Similarly, an extreme loss of determinacy in florets was also observed in *zag1-mum1; drl1-R; drl2-M* triple mutants (Supplementary Fig. 2e). The synergistic genetic interactions between *drl1* and *drl2* mutant and natural variant alleles in florets were dose-sensitive, consistent with dosage effects observed for leaf traits^23^. Varying the dosage of *drl2-M* in an F_2_ population resulted in florets of *drl1-R; drl2-M/+* plants being intermediate in severity between *drl1-R* homozygotes and *drl1-R; drl2-M* double homozygotes (Fig. 1c). In support of these observations, classification of DRL1 as a “regulator” in a gene regulatory network (GRN) from integrated transcriptome, proteome and phosphoproteome datasets across a developmental atlas for maize^24^ revealed high-confidence edges with *drl1* and *drl2* “targets,” indicative of putative auto- and cross-regulation activities, respectively (Supplementary Fig. 4). Collectively, these observations suggest that the *drl* loci regulate pistillate floret development and FM determinacy in a dose-dependent manner.

**Figure 3.**
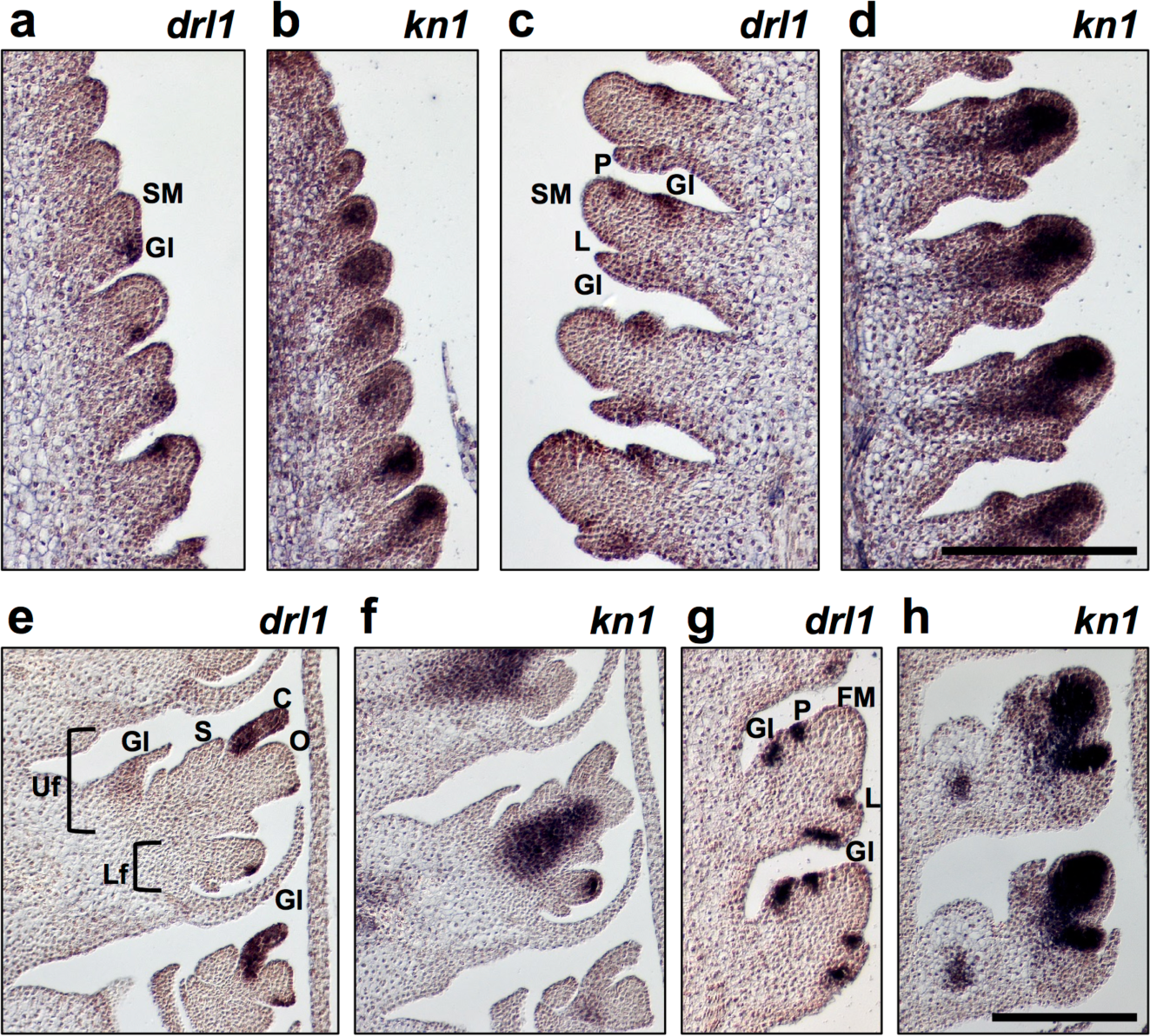
*drl* transcript accumulation in normal pistillate and staminate florets. **a-d,** Longitudinal sections through developing B73 ears hybridized with antisense RNA probes to *drl1* (**a,c**) or *kn1* (**b,d**). **e,f,** Longitudinal sections through developing B73 pistillate florets hybridized with antisense RNA probes to *drl1* (**e**) or *kn1* (**f**). **g,h,** Longitudinal sections through developing B73 staminate florets hybridized with antisense RNA probes to *drl1* (**g**) or *kn1* (**h**). SM, spikelet meristem; Gl, glume primordium; L, lemma primordium; P, palea primordium; Uf, upper floret; Lf, lower floret; C, carpel primordium; O, ovule primordium; S, stamen primordium. Scale bars, 200 µm.

**Figure 4.**
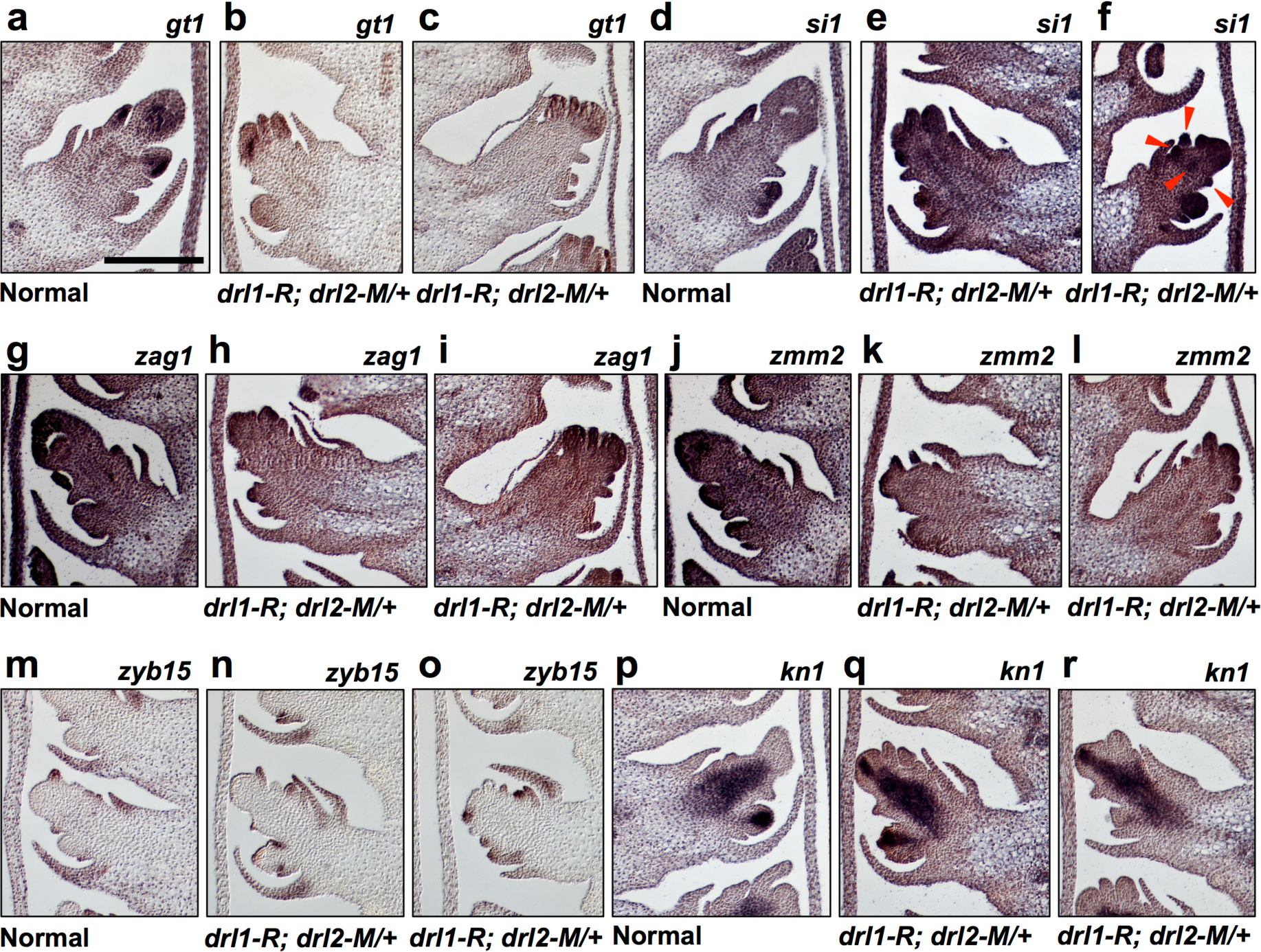
Transcript accumulation of marker genes in *drl1-R; drl2-M/+* pistillate florets. **a-r,** Longitudinal sections through late-stage pistillate florets from normal (**a,d,g,j,m,p**) and *drl1-R; drl2-M/+* (**b,c,e,f,h,i,k,l,n,o,q,r**) siblings hybridized with antisense RNA probes to *gt1* (**a-c**), *si1* (**d-f**), *zag1* (**g-i**), *zmm2* (**j-l**), *zyb15* (**m-o**) and *kn1* (**p-r**). Arrowhead points to ectopic accumulation of *si1* transcripts in *drl1-R; drl2-M/+* florets. Scale bar, 200 µm.

In the *drl1* mutant tassel, an ectopic stamen appeared periodically in the UF of sessile (3.14 ± 0.06) and pedicellate (3.05 ± 0.04) spikelets (Fig. 1e,g). Histological examination of mature *drl1* mutant spikelets revealed the infrequent extra stamens originated internal to the normally placed and numbered lemma and palea in the outer whorl (Fig. 1f). We did not observe a nucellus in *drl1* mutant staminate florets. The minor degree of FM indeterminacy in *drl1* mutants was enhanced in *drl1; drl2* double mutants where stamen number increased in both the UF and lower floret (LF) of sessile (4.49 ± 0.09, *P* = 1.0 × 10^−19^; 3.41 ± 0.08, *P* = 4.6 × 10^−6^,respectively) and pedicellate (3.92 ± 0.11, *P* = 3.0 × 10^−10^; 3.19 ± 0.08, *P* = 0.015, respectively) spikelets (Fig. 1e-g). Differences were significant between stamen number in the UF and LF within sessile and within pedicellate spikelets (*P* < 10^−6^), and for UFs, between sessile and pedicellate spikelets (*P* < 10^−3^) (Fig. 1g). These data suggest that the *drl* genes participate differentially in determinacy pathways of upper and lower staminate FMs, and that ectopic stamens originate from sustained activity of the mutant FM.

*drl1; drl2* double mutant florets displayed phenotypes that were not observed in *drl1* single mutants. An ectopic primordium with lodicule-like cellular morphology and vascularization was observed occasionally in position of a presumptive, suppressed adaxial-medial lodicule in the UF (Fig. 1f, right panel, arrowhead), indicating a possible role for *drl* gene products in imposing zygomorphy^20,25^. We also observed macrohair-like structures along the apical ridge of *drl1-R; drl2-M* supernumerary anthers (Supplementary Fig. 5). Macrohair production is generally limited to the adaxial epidermis of the adult leaf blade and is frequently used as a morphological marker for leaf polarity^26^. Though ectopic structures were infrequent, they lacked the multicellular bases of leaf blade macrohairs^27^ and were consistently associated with supernumerary anthers with altered morphology. Such amorphic anthers had aberrant theca that lacked pollen sacs and were often fused to morphologically normal anthers.

**Figure 5.**
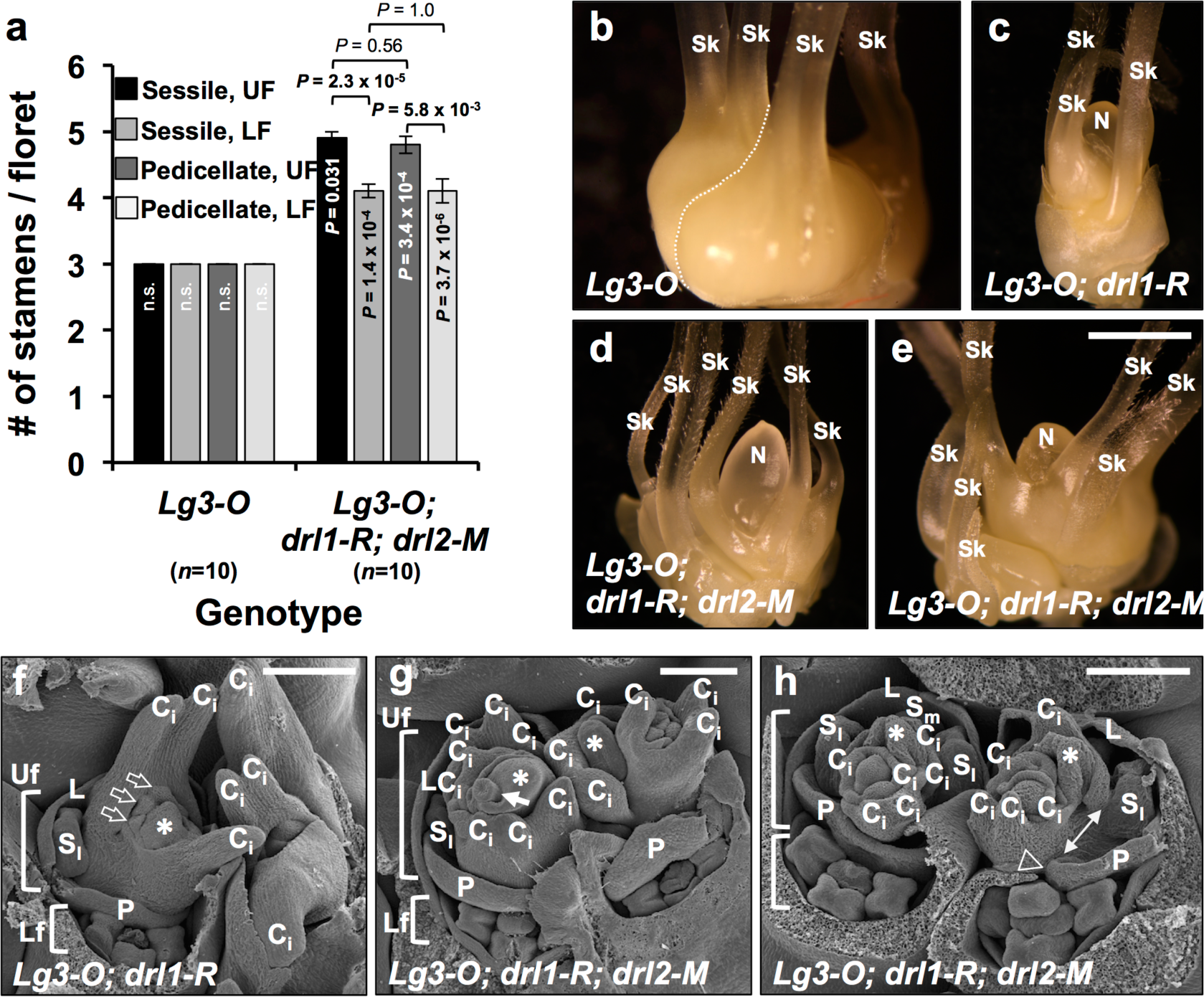
Genetic interaction between *drl1, drl2* and *Lg3* mutants. **a,** Quantification of stamens in *Lg3-O* and *Lg3-O; drl1-R; drl2-M* florets. Mean ±SEM, *P*-values based on two-tailed Student’s *t* tests compared to *drl1-R; drl2-M* double mutants in Fig. 1g; *n*, sample size. **b-e,** Dissected pistillate spikelet of *Lg3-O* (**b**), *Lg3-O; drl1-R* (**c**) and *Lg3-O; drl1-R; drl2-M* (**d,e**). **f-h,** Scanning electron micrographs of late-staged pistillate florets from *Lg3-O; drl1-R* (**f**) and *Lg3-O; drl1-R; drl2-M* (**g,h**). Glumes were removed manually to expose the upper and lower florets. N, nucellus; Sk, silk; Uf, upper floret; Lf, lower floret; L, lemma primordium; P, palea primordium; S_l_, lateral stamen primordium, S_m_, medial stamen primordium, C_i_, indeterminate carpel primordium; C_d_, determinate carpel primordium. Asterisks (**f-h**) mark ectopic primordium. Open arrows (**f**) point to initiation of three closely-spaced carpel primordia. Closed arrow (**g**) points to reduced FM. Double arrow (**h**) marks aberrant spacing between palea axil and lateral stamen primordium. Arrowhead (**h**) points to palea involution. Scale bars, 1 mm (**b-e**) and 200 µm (**f-h**).

### *drl1* and *drl2* impose FM determinacy

We tracked the developmental basis of *drl1* and *drl1; drl2* mutant phenotypes in mid- and later-staged pistillate florets with scanning electron microscopy (SEM). Prior to sex determination, the inner whorl of normal UFs consists of a medial-adaxial C_d_ primordium (determinate carpel) and two lateral-adaxial C_i_ primordia (indeterminate carpels), all of which are connately fused (Fig. 2a). This gynoecial whorl is flanked by a whorl of three pre-degenerate stamen primordia. The LF lags in development, here with a FM and recently initiated stamen primordia (Fig. 2a). In mid-staged UFs of *drl1* mutants, the medial-adaxial C_d_ failed to initiate (Fig. 2b), resulting in the single protruding nucellus observed in mature *drl1* mutant florets (Fig. 1d). *drl1* mutant UFs had extra whorls of lateral C_i_ (Fig. 2c); however, shifts in phyllotaxis between each extra whorl complicated assigning ab- or adaxial orientation relative to the palea axil. The medial-adaxial C_d_ was similarly suppressed in mid-staged UFs of *drl1; drl2* double mutants, yet they displayed multiple whorls of lateral C_i_ indicating prolonged FM activity (Fig. 2d-f). Additionally, ectopic primordia, interpreted to each be unexpanded nucellus based on position, initiated in the axil of each C_i_ whorl of *drl1; drl2* double mutants (Fig. 2d-f, asterisks), and likely accounts for the multiple, expanded nucelli observed in mature *drl1; drl2* double mutant florets (Fig. 1d).

In normal later-staged UFs, lateral-adaxial C_i_ primordia appear as bi-keeled (paired) and elongate, while the reduced medial-adaxial C_d_ envelops the ovule (Fig. 2g). Multiple whorls of paired lateral-adaxial C_i_ primordia were obvious in similarly staged *drl1* mutant UFs (Fig. 2h), whereas tri-keeled lateral-adaxial C_i_ primordia observed in later-staged UFs of *drl1; drl2* double mutants indicated the presence of an extra, intra-whorl fused or partially fused C_i_ primordium (Fig. 2i). Additionally, we often detected involution of the palea along the medial axis in *drl1; drl2* double mutant florets (Fig. 2e,i, arrowheads), which may indicate crowding within the inner whorl of the floret. Taken together, these observations document that the *drl* genes are required for proper elaboration of pistillate florets, including C_d_ initiation, and to impose FM determinacy.

### *drl* genes are expressed solely in lateral primordia and dynamically throughout inflorescence development

The *drl* genes are required for carpel development and to impose FM determinacy (Figs 1 and 2). To examine the temporal and spatial patterns of *drl* transcript accumulation during inflorescence and floret development, we performed RNA *in situ* hybridization. In median longitudinal sections of the developing ear, *drl1* transcripts were detected in the IM periphery, which spatially correspond to cryptic bract anlagen^28^ (Supplementary Fig. 6a). *drl1* transcripts continued to accumulate in outer glume primordia (Fig. 3a), but not in the SM, as marked by accumulation of *knotted1* (*kn1*) transcripts^29^ (Fig. 3b). The accumulation pattern of *drl1* transcripts persisted in later-staged SMs where they were detected in lemma and palea primordia (Fig. 3c,d), expression patterns that were also observed for *drl2* (Supplementary Fig. 6b,c). In more advanced pistillate florets, *drl1* transcripts accumulated in carpel primordia that had initiated in the UF and LF, but not in the central presumptive ovule primordium of either floret (Fig. 3e,f). In developing staminate florets, *drl1* transcripts accumulated in a similar pattern in lateral primordia that were initiated by the FM (Fig. 3g), but not within the FM (Fig. 3h). *drl* expression dynamics across developing inflorescences were supported using publicly available RNA-seq data (Supplementary Fig. 6d) (www.maizeinflorescence.org). To summarize, the *drl* genes are expressed in cryptic bracts, in lateral organ primordia initiated by the SM (glumes), and in primordia of outer- (lemma and palea) and inner- (carpels) whorl organs initiated by the FM. *drl* expression in carpel primordia correlates with the organs that were altered in *drl1* and *drl1; drl2* mutant florets. The indeterminate FMs observed in *drl* mutant florets are best explained by misregulation of FM activity, yet *drl* expression is limited to organs derived from the meristems and is excluded from the meristem. These points strongly suggest that *drl* regulates meristem activity via a non-cell autonomous mechanism. Consistent with this hypothesis, *drl1* and *drl1; drl2* mutants also display a dose-dependent reduction in vegetative shoot apical (SAM) meristem size even though the *drl* genes are expressed in leaf primordia and not in the SAM proper^23^.

### Expression patterns of floret markers in *drl1; drl2* mutants indicate sustained FM activity

We performed RNA *in situ* hybridization with floret marker genes to understand the basis for the indeterminacy and potential organ identity shifts in pistillate florets of *drl* mutants. We used *drl1-R; drl2-M/+* individuals (Fig. 1c) for these analyses. The maize *grassy tillers1* (*gt1*) gene encodes a homeodomain leucine zipper transcription factor that is required to repress carpel growth in staminate florets^30^. In developing tassel florets, *gt1* is expressed in the central gynoecium^30^. We examined longitudinal sections through later-staged, normal pistillate florets, where *gt1* transcripts accumulated in the central gynoecium and in stamen primordia (Fig. 4a).

In contrast, in *drl* mutant UFs, *gt1* transcripts accumulated broadly throughout the presumptive ovule primordium and to a lesser degree throughout the LFM (Fig. 4b,c). Accordingly, GRN subnetwork analysis^24^ revealed a high-confidence edge score for *gt1* as a putative target of DRL1 (Supplementary Fig. 4). The maize *APETALA3/DEFICIENS* ortholog, *silky1* (*si1*), is required for lodicule and stamen identity, and in staminate and pistillate florets is expressed in developing lodicule and stamen primordia^31,32^, where *si1* transcripts accumulate asymmetrically^25^. As expected, we found that *si1* transcripts accumulated in degenerating stamen primordia in the normal pistillate UF and in the LF, whereas in *drl1-R; drl2-M/+* the accumulation pattern was broader and included ectopic primordia in the UF and strong expression throughout the LF (Fig. 4d-f). The *zag1* gene regulates FM determinacy with a lesser role in promoting carpel identity; currently, functional analyses have not been reported for its duplicate factor *zea mays mads2* (*zmm2*)^7^. *zag1* and *zmm2* expression domains overlap largely throughout the development of pistillate florets, where they mark the FM as well as stamen and carpel primordia^32,33^.

Expression patterns for *zag1* and *zmm2* differ in developing staminate florets^25^. We observed that *zag1* transcripts accumulate in stamen and carpel primordia of *drl1-R; drl2-M/+* late-staged pistillate UFs and in the LF, whereas in normal florets, accumulation was largely confined to the incipient silk (Fig. 4g-i). Transcripts of *zmm2* accumulated in a similar pattern in normal and *drl1-R; drl2-M/+* developing pistillate florets (Fig. 4j-l). Interestingly, GRN subnetwork analysis^24^ revealed high-confidence edge scores for 23 annotated MADS-box genes that included *si1*, *zag1* and *zmm2* as putative DRL1 targets; GRN analysis^24^ also predicted ZAG1 as a putative regulator of *drl1* and *drl2* genes (Supplementary Fig. 4). The *FILAMENTOUS FLOWER* homolog, *zea yabby15*/*yabby8* (*zyb15*/*yab8*)^23,26^ is expressed during inflorescence development in cryptic bract primordia^28^. We observed *zyb15* transcript accumulation in glume, lemma, palea and carpel primordia, but not in the FM or in stamen primordia, for both normal and *drl1-R; drl2-M/+* developing pistillate UF and LF (Fig. 4m-o). Interestingly, *zyb15* transcript accumulation persisted longer in glume primordia compared to *drl1* accumulation (Fig. 4m cf. 3e). The *kn1* gene is expressed in meristematic cells and is downregulated in cells recruited to form a lateral domain on the flank of the meristem and in lateral organ primordia^29^. We observed that *kn1* transcripts were absent from normal later-staged pistillate UFs that had undergone terminal differentiation to an ovule primodium, whereas in similarly staged *drl1* mutant UFs, *kn1* transcript accumulation persisted throughout the gynoecial axis, demonstrating the *drl1; drl2* FMs are indeterminate (Fig. 4p-r). In summary, the expression patterns of marker genes in developing *drl1-R; drl2-M/+* pistillate florets shed light on the genetic basis of indeterminate floret growth in *drl1-R; drl2-M* double mutants. We observed notable shifts from normal patterns of transcript accumulation for *gt1* where transcripts accumulated more broadly throughout the indeterminate pistillate floral axis (Fig. 4b,c), for *si1* where the pattern of ectopic accumulation may indicate that in *drl1-R; drl2-M* mutants, extra stamen primordia initiate in pistillate UFs or lodicule primordia are de-repressed (Fig. 4f), and for *kn1*, where prolonged transcript accumulation marked the *drl1-R; drl2-M/+* indeterminate floral axis (Fig. 4q,r).

### *drl* genes interact with the pistil abortion pathway

Maize is monoecious with unisexual florets borne on the terminal tassel and lateral ear. Tassels resemble ears in the *tasselseed1* (*ts1*) mutant in that feminization of tassel spikelets reduces glume growth, arrests stamen growth, and permits development and growth of a fertile, determinant gynoecial whorl^34,35^. The *ts1* gene encodes a lipoxygenase that may act during the biosynthesis of jasmonic acid^36^. Laudencia-Chingcuanco and Hake (2002) reported an occasional determinant nucellus in staminate florets of *ifa1* mutants. In *drl1; drl2* mutants, we observed extra anthers indicative of reduced FM determinacy in staminate florets; however, with multiple expanded nucelli, loss of FM determinacy was more evident in pistillate florets (Fig. 1). To test for an interaction between pathways that regulate FM determinacy and pistil abortion, we examined F_2_ progeny between *drl1-R; drl2-M* double mutants and the *ts1-Alex* mutant. Tassel florets of *ts1-Alex; drl1-R; drl2-M* triple mutants were feminized like *ts1-Alex*, but additionally showed many hallmarks of *drl1; drl2* double mutant pistillate (ear) florets: reduced, unfused carpels and multiple ectopic nucelli in the UF and LF of the tassel spikelet (Supplementary Fig. 7). Thus, *ts1-Alex* markedly enhanced the indeterminacy of *drl1-R; drl2-M* mutant tassel florets (Supplementary Fig. 7b,c). Interestingly, in triple mutants, we consistently observed nucelli in tassel florets were less expanded than nucelli in ear florets, indicating that additional factors may contribute differentially to nucellus growth in ear versus tassel florets. Taken together, the degree of FM indeterminacy differs in *drl1; drl2* double mutant staminate and pistillate florets, but that difference diminishes substantially when pistil abortion is suppressed in tassel florets via the *ts1-Alex* mutation. These observations suggest that the *drl* genes control stem cell proliferation perhaps by integrating hormonal cues such as from lateral primordia, and that specific integration mechanisms in staminate and pistillate florets may respond to different hormone levels or sensitivities^34,35^.

### *drl* and *knox* pathways interact genetically to regulate FM activity

Recessive loss-of-function mutations in *kn1*^37^, or dominant gain-of-function mutations in the class I *kn1-like homeobox* (*knox*) genes *Gnarley1*^38^ or *Liguleless3-O* (*Lg3-O*) (this study, Fig. 5) can display ectopic carpels in pistillate florets, with no reported changes in stamen number in tassel florets. Similarly, pistillate florets of recessive loss-of-function mutations in *rough sheath2* (*rs2*), a negative regulator of *knox* genes^39-41^, occasionally display multiple silks, while stamen number remains normal in tassel florets (Supplementary Fig. 8)^39^. To determine whether the *drl* genes function in the *knox* pathway or pathway(s) that regulate KNOX activity, we examined the genetic interactions between *drl1; drl2* and *Lg3-O* or *rs2-R* mutants.

Stamen number in *Lg3-O* mutants did not deviate from normal (Fig. 5a cf. 1g). However, we observed significant enhancement of ectopic stamens in UF and LF of both sessile and pedicellate spikelets of *Lg3-O; drl1-R; drl2-M* triple mutants compared to *drl1-R; drl2-M* double mutants (Fig. 5a cf. 1g). Similar to what we observed in *drl1-R; drl2-M* double mutants, for *Lg3-O; drl1-R; drl2-M* triple mutants the differences in stamen number between UF and LF were highly significant within sessile and pedicellate spikelets (*P* < 10^−2^) (Fig. 5a). In pistillate florets of *Lg3-O* mutants, we found supernumerary carpels that formed partial, extra pistils, which occasionally did not contain an ovule (Fig. 5b), an observation not reported previously^42,43^.

Dissected pistillate florets from *Lg3-O; drl1-R* double and *Lg3-O; drl1-R; drl2-M* triple mutants revealed ectopic, unfused carpels that failed to enclose a central nucellus (Fig. 5c-e).

Interestingly, the multiple nucelli that we observed in triple mutants were reduced in their growth compared to those in *drl1-R; drl2-M* double mutants (Fig. 5d,e cf. 1d). SEM analysis of developing *Lg3-O; drl1-R* double mutant pistillate florets revealed carpel elongation from both C_i_ and C_d_ gynoecial ridges (Fig. 5f), whereas in *Lg3-O; drl1-R; drl2-M* triple mutants, multiple whorls of C_i_ were apparent around the central FM axis (Fig. 5g,h), which led to consumption of the FM in some florets (Fig. 5g). In some *Lg3-O; drl1-R; drl2-M* triple mutants, the stamen whorl was aberrantly arranged within the palea axil (Fig. 5h). Collectively, these data indicate that misexpression of *lg3* enhances aspects of the *drl1-R; drl2-M* staminate and pistillate floret phenotypes. However, because *Lg3-O* is characterized by a gain-of-function mutation, an exact functional relationship between the *drl* genes and *lg3* is difficult to interpret. The floret phenotypes in *Lg3-O* and *drl* higher-order mutants could indicate that these genes operate in parallel pathways or in the same pathway.

We generated *rs2-R; drl1-R; drl2-M* triple mutants to ask if the *drl* genes intersect with, or operate in parallel to, the *ASYMMETRIC LEAVES1/PHANTASTICA* ortholog *rs2*, which is required to repress multiple *knox* gene activities, including *lg3*, at sites of organ initiation^39-41^. In pistillate florets, *rs2* transcripts accumulate in lateral organs initiated by the SM and FM^40^ in domains comparable with *drl* expression (Fig. 3). We found that stamen number in *rs2-R* did not deviate from normal and observed significant differences in ectopic stamen number between UF and LF within both sessile and pedicellate spikelets of *rs2-R; drl1-R; drl2-M* triple mutants (*P* < 10^−4^; Supplementary Fig. 8a). The differences were greater than those observed in *drl1-R; drl2-M* double mutants, implying enhancement by *rs2-R*. Interestingly, the arrangement of inner whorl organs such as lodicules, stamens and ectopic stamen-like structures was irregular in staminate florets of *rs2-R; drl1-R; drl2-M* triple mutants, and lodicules tended to be amorphic (Supplementary Fig. 8b). Ectopic stamens and stamen-like structures often derived from a central site in the inner whorl and were frequently fused along their filaments. Dissected *rs2-R* pistillate florets can have multiple silks with fused carpels (Supplementary Fig. 8c)^39^, whereas *rs2-R; drl1-R; drl2-M* pistillate florets displayed extreme indeterminacy, with numerous carpelloid-like structures that appeared to derive from multiple origins in the UF (Supplementary Fig. 8d). These carpelloid-like structures were reduced, fleshy, and did not appear to surround a central nucellus, which was conspicuously absent compared to *drl1-R; drl2-M* pistillate florets (Fig. 1d). Taken together, these results indicate the *drl* and *rs2* genes likely converge on shared targets to regulate staminate and pistillate FM determinacy.

## Discussion

Floret architecture in the cereals is a major component of yield. Elegant dissections of Arabidopsis *CRC* and rice *DL* gene function have contributed to our understanding of how these *YABBY* genes regulate floral development across species with perfect, bisexual flowers^12,13,14,44^. Our results demonstrate that the maize *drl* genes are required to impose FM determinacy and for proper elaboration of inner whorl organs of staminate and pistillate florets, indicating critical roles for *drl* genes in regulating stem cell homeostasis and organ growth in dimorphic, unisexual florets (Figs 1 and 2). We hypothesize the *drl* gene products function non-cell autonomously in or through pathways that signal from lateral primordia, through boundary domains, to regulate developmental programs of the FM. Our findings suggest that non-cell autonomous function of *drl* interacts differentially with the distinct developmental potentials of staminate UFMs, LFMs and pistillate UFMs (Figs 1-3 and 6). UFs and LFs differentially express key regulators^45,46^, potentiate differential effects of developmental regulators^10^, and derive from slightly different developmental trajectories of the SM^47^. Perhaps akin to the maize *ZmFON2-LIKE CLE PROTEIN1* (*FCP1*)*-FASCIATED EAR3* primordia-to-meristem feedback circuit^48^, a feedback signaling system from SM- and FM-derived lateral primordia involving the *drl* gene products could provide vital control of stem cell proliferation by integrating hormonal or metabolic cues from incipient and emerging primordia. With some 48 *CLE* genes currently reported in maize^49^, it is tempting to speculate that differential interactions between *drl* and *CLE* genes and/or gene products could provide non-cell autonomous regulation of FM activity from lateral floral primordia. Indeed, GRN subnetwork analysis^24^ uncovered high-confidence edge scores for *CLE-FCP1* genes as putative DRL1 targets (Supplementary Fig. 4). In broader context, a recent report in *Caenorhabditis elegans* underscores how non-cell autonomous signaling from somatic to adjacent germline tissue regulates stem cell proliferation in the germline^50^, implying a common theme in development used to control stem cell proliferation.

## Methods

### Genetic stocks and plant growth

Maize plants were grown in the field or in the greenhouse. The *drl1* and *drl2* alleles used in this study were described previously^23^. *drl* alleles were backcrossed to A619, B73, Mo17 and W22 inbred lines at least four times. The *Lg3-O* (B73-5) allele was obtained from Michael Muszynski (Iowa State University). The *rs2-R* (Mo17-many) and *ts1-Alex* (Mo17-4) alleles were obtained from Erik Vollbrecht (Iowa State University). The *ifa1* (B73-4) allele was obtained from Sarah Hake (UC-Berkeley). The *zag1-mum1* (B73-many) allele was obtained from David Jackson (Cold Spring Harbor Laboratory). For quantitative phenotyping, sample sizes per genotype are indicated throughout the manuscript, along with mean ± s.e.m. presented with significance calculated using two-tailed Student’s *t* tests. All experiments were performed with two or three independent biological replicates.

### Genetic interaction analysis

Higher-order mutants were generated using the *drl1-R* and *drl2-M* alleles and *ifa1*, *zag1-mum1*, *ts1-Alex*, *Lg3-O*, or *rs2-R* alleles. The F_1_ progeny from these crosses were grown to maturity and self-pollinated or backcrossed. The F_2_ or BC_1_ progeny were grown to maturity and screened for the *drl1-R* and *drl2-M* alleles by genotype and for the tester mutant alleles by phenotype, except for *Lg3-O* (Lg3_13-CGTCCATTTCCCATCCCCAA and Lg3_6-CCTTGCGGCACTCGATGTA), *rs2-R* (rs2_F3-CGCATTATGAGGTGTGGTGG and rs2_R1-CTCCATCTCCAGCTGCTGC) and *zag1-mum1* (zag1_F2-GGAATCTGCTAGGCTGAGGC and zag1_R2-GGTCGTTGAAGTCTTTCCGG) alleles, which were genotyped. Genotyping primers for *ifa1*, *drl1-R* and *drl2-M*, as well as DNA isolation and PCR conditions were described previously^23^.

### Histology

Toluidine blue O (TBO) (Sigma) staining was performed on mature spikelets. Briefly, TBO was dissolved in 1% sodium borate (w/v) to make a 1% stock solution (w/v). A 0.5% TBO staining solution was made immediately before use by diluting the stock solution with 1% sodium borate. Microtome sections of 10 µm, adhered to a microscope slide, were deparaffinized in Histo-Clear (National Diagnostics) (2 times, 10 min. each). Slides were passed through a graded ethanol series toward hydration, 1 minute each (100%, 100, 95, 95, 70, 50, distilled water) and stained in 0.5% TBO staining solution for 3 minutes. Slides were then passed through a graded series toward dehydration, 30 seconds each (50%, 70, 95, 95, 100, 100) and Histo-Clear (3 times, 5 min. each). Slides were coverslip mounted with Permount (Fisher).

### Scanning Electron Microscopy

Field-grown ears 10 mm in length were fixed with 2% paraformaldehyde and 2% glutaraldehyde in cacodylate buffer (0.1 M) at pH 7.2 for at least 24 hours / 4 ˚ C. After fixation, samples were rinsed 3 times (15 min. each) in cacodylate buffer (0.1 M). Then samples were post-fixed in 1% osmium tetroxide in cacodylate buffer (0.1 M) for 1 hour. After several washes with deionized water, samples were dehydrated through a graded ethanol series (25%, 50, 70, 85, 95, 100) 2 changes each for 15 minutes. Samples were critical point dried using a Denton Vacuum, Inc. Drying Apparatus, Model DCP-1 (Denton Vacuum, Moorestown, NJ). Dried materials were mounted on aluminum stubs with double-sided tape and colloidal silver paint and sputter coated with gold-palladium with a Denton Desk II Sputter Coater (Denton Vacuum, Inc. Moorestown, NJ). Images were captured using a JEOL JSM-5800LV scanning electron microscope at 10 kV (JEOL = Japan Electronic Optics Laboratory, Peabody, MA).

### RNA *in situ* hybridization

Field-grown 10 mm maize ears were fixed overnight at 4 ˚ C in 3.7% FAA. Samples were dehydrated through a graded ethanol series (50%, 70, 85, 95, 100) each 1 hour, with 2 changes in 100% ethanol. Samples were then passed through a graded Histo-Clear (National Diagnostics) series (3:1, 1:1, 1:3 ethanol: Histo-Clear) with 3 changes in 100% Histo-Clear; all changes were 1 hour each. Samples were then embedded in Paraplast®Plus (McCormick Scientific), sectioned, and hybridized as described previously^23^. Hybridizations were performed using antisense digoxygenin-labeled RNA probes: *drl1*^23^, *drl2*^23^, *kn1*^29^, *gt1*^30^, *si1*^25^, *zag1*^25^, *zmm2*^25^, and *zyb15/yab8* (JS137-CGATCTCTACGCCGCAGC and JS138-GCAGACATACGCAAACATGGG).

### Accession numbers

Maize: *drl1*, GRMZM2G088309; *drl2*, GRMZM2G102218; *gt1*, GRMZM2G005624; *kn1*, GRMZM2G017087; *lg3*, GRMZM2G087741; *rs2*, GRMZM2G403620; *si1*, GRMZM2G139073; *ts1*, GRMZM2G104843; *zyb15/yab8*, GRMZM2G529859; *zag1*, GRMZM2G052890; *zmm2*, GRMZM2G152862.

## Acknowledgements

We thank Harry Horner and Tracey Stewart at the Iowa State University Bessey Microscopy Facility for assistance with scanning electron microscopy and Pete Lelonek for plant care. Many thanks to Clint Whipple for generously sharing *gt1*, *si1*, *zag1* and *zmm2* probes for RNA *in situ* hybridization, and to Beth Thompson for discussions on *ifa1*. We thank Justin Walley for helpful discussions on incorporating GRN analysis. We are grateful for the many former undergraduate students, especially Sarah Briggs, Emery Peyton and Charlie Beeler, for their help in our summer genetics nurseries. We also thank Jim Cahill for the *Lg3-O* genotyping assay. Many thanks to Erin Irish and Erica Unger-Wallace for insightful discussions and comments on the manuscript. This work was supported by the National Science Foundation (IOS-1238202).

## Author contributions

J.S. designed and performed the experiments, and analyzed the data with guidance from E.V.

J.S. wrote the article with edits from E.V.

## Supplementary information

**Figure S1.**
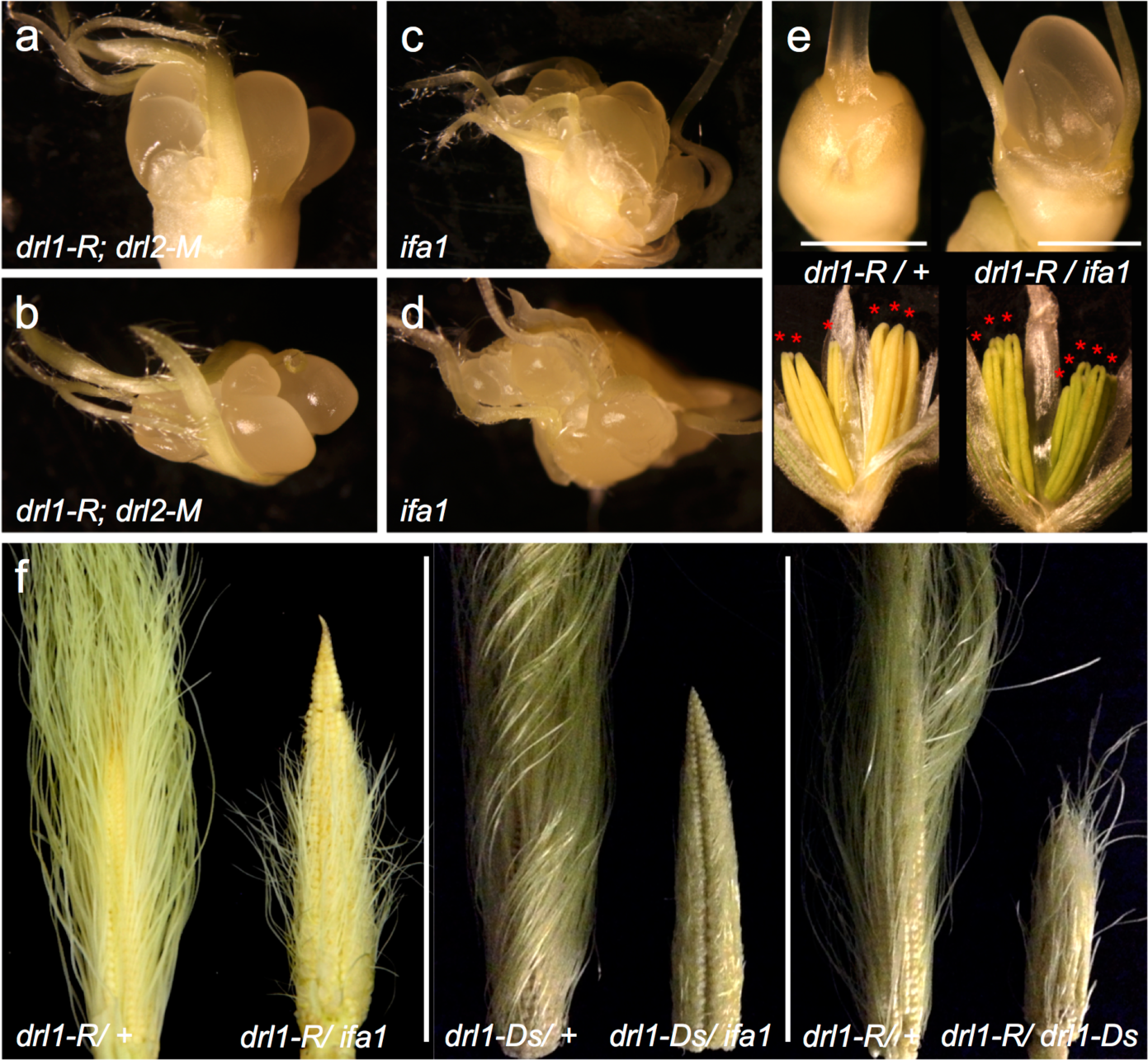
Complementation tests for *drl1* alleles. **a-d,** Dissected pistillate florets showing multiple nucelli in *drl1-R; drl2-M* (**a,b)** and in *drl1-ifa1* (**c,d**). (**e**) Dissected pistillate and staminate florets of F_1_ progeny of *drl1-R/+* heterozygous and *drl1-R/ifa1* trans-heterozygous siblings. **(f)** Mature ears of *drl1-R/+* sibling and *drl1-R/ifa1* (left panel); *drl1-Ds/+* sibling and *drl1-Ds/ifa1* (middle panel); and *drl1-R/+* sibling and *drl1-R/drl1-Ds* (right panel). Asterisks mark mature anthers. Scale bar, 2 mm.

**Figure S2.**
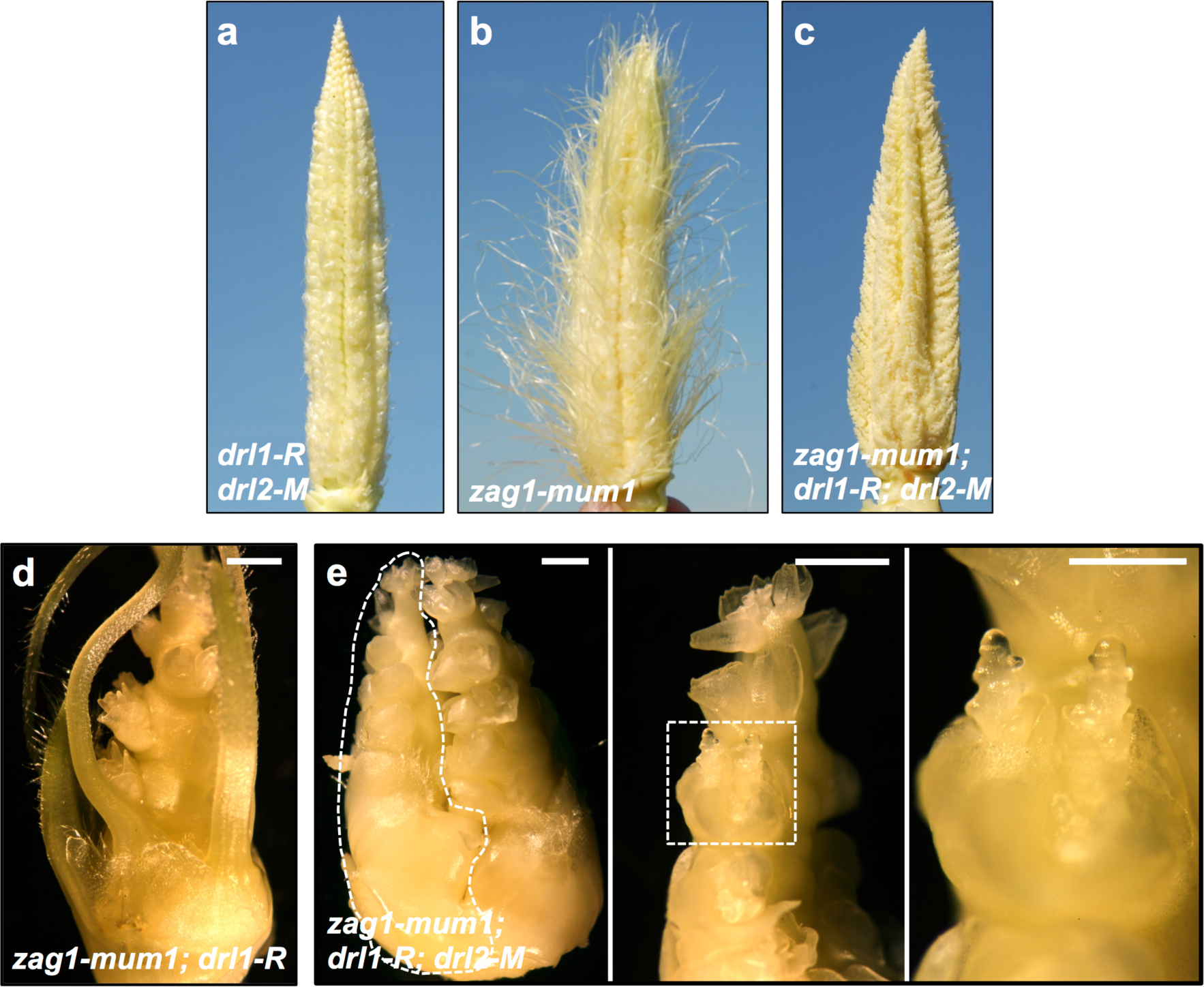
Genetic interaction between *drl1, drl2* and *zag1* mutants. **a-c,** Mature ears of *drl1-R; drl2-M* (**a**), *zag1-mum1* (**b**) and *zag1-mum1; drl1-R; drl2-M* (**c**). **d,e,** Dissected pistillate spikelet of *zag1-mum1; drl1-R* (**d**) and *zag1-mum1; drl1-R; drl2-M* (**e**). **e,** Branch-like floret outlined (left) shown at high-magnification (middle) with boxed region at higher-magnification (right). Scale bars, 1 mm.

**Figure S3.**
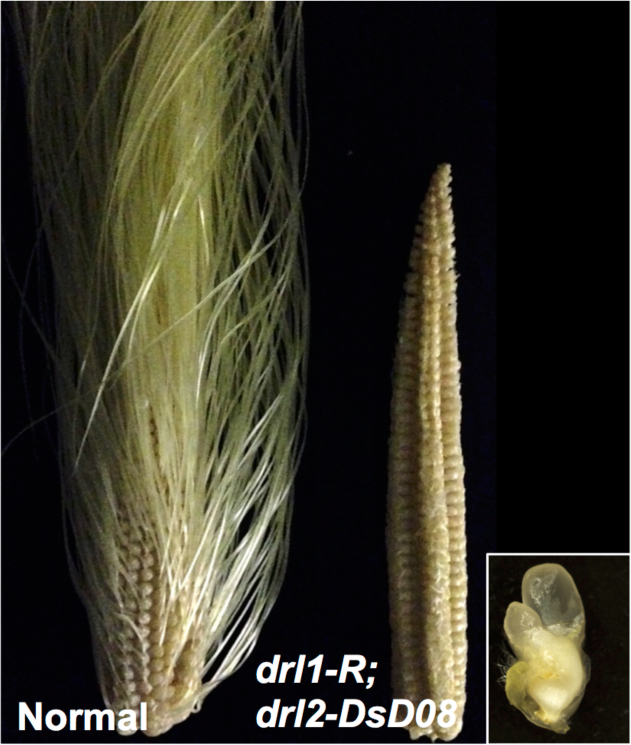
Mature ear and floret of *drl1-R; drl2-DsD08*. Mature ears of normal sibling and *drl1-R; drl2-DsD08* in the W22 (non-enhancing) inbred background. Inset, dissected *drl1-R; drl2-DsD08* floret showing multiple nucelli.

**Figure S4.**
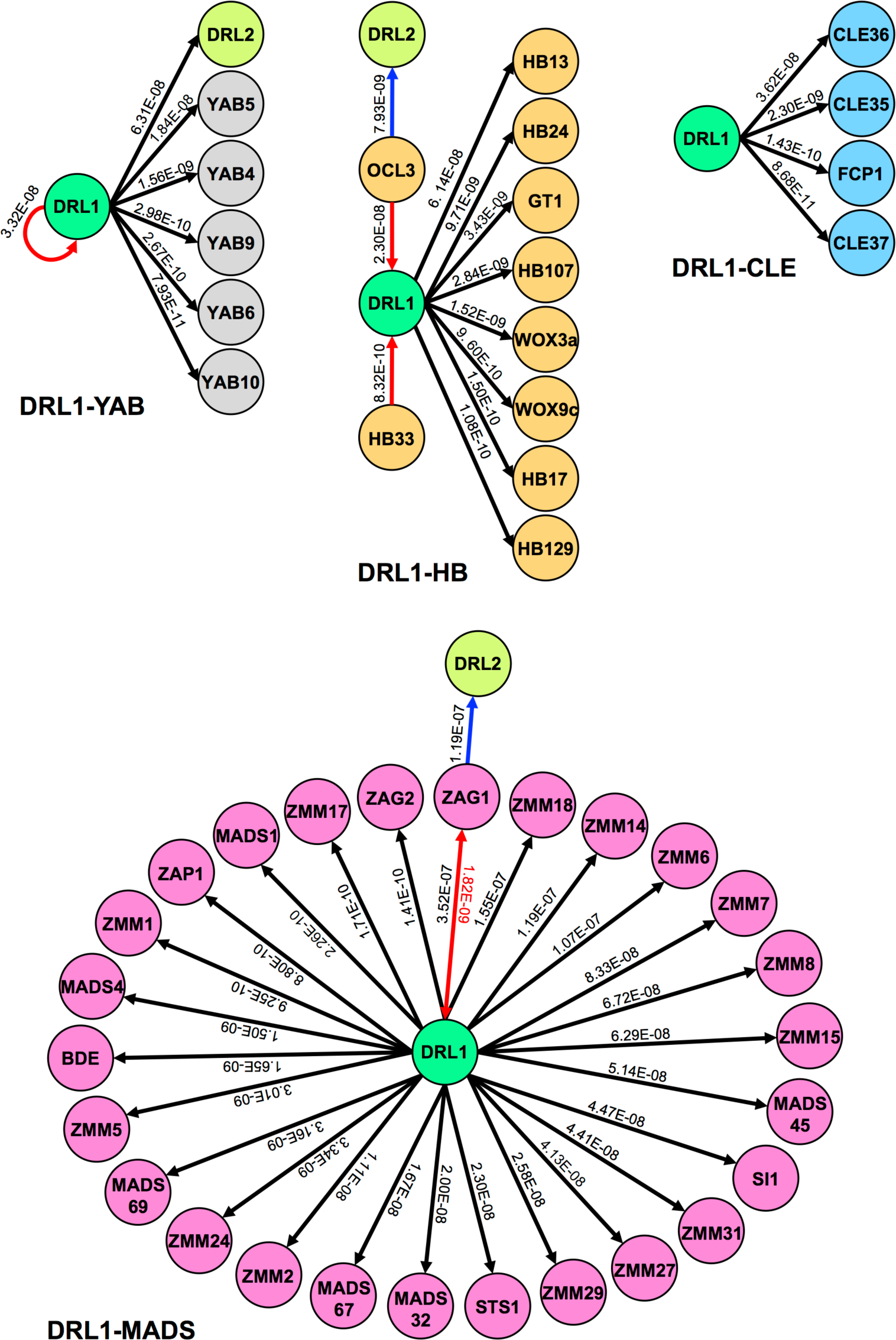
Unsupervised GRN subnetworks24 centered on DRL1. GRN subnetworks for DRL1-YABBY (DRL1-YAB), DRL1-HOMEODOMAIN (DRL1-HB), DRL1-CLV EMBRYO SURROUNDING REGION (DRL1-CLE) and DRL1-MADS BOX (DRL1-MADS). Gene annotations for DRL1-YAB, DRL1-HB and DRL1-MADS subnetworks were taken from www.grassius.org and DRL1-CLE from ref. 49. Regulator-target and relationship of interaction are depicted by arrows representing high-confidence edges with edge score value above each edge.

**Figure S5.**
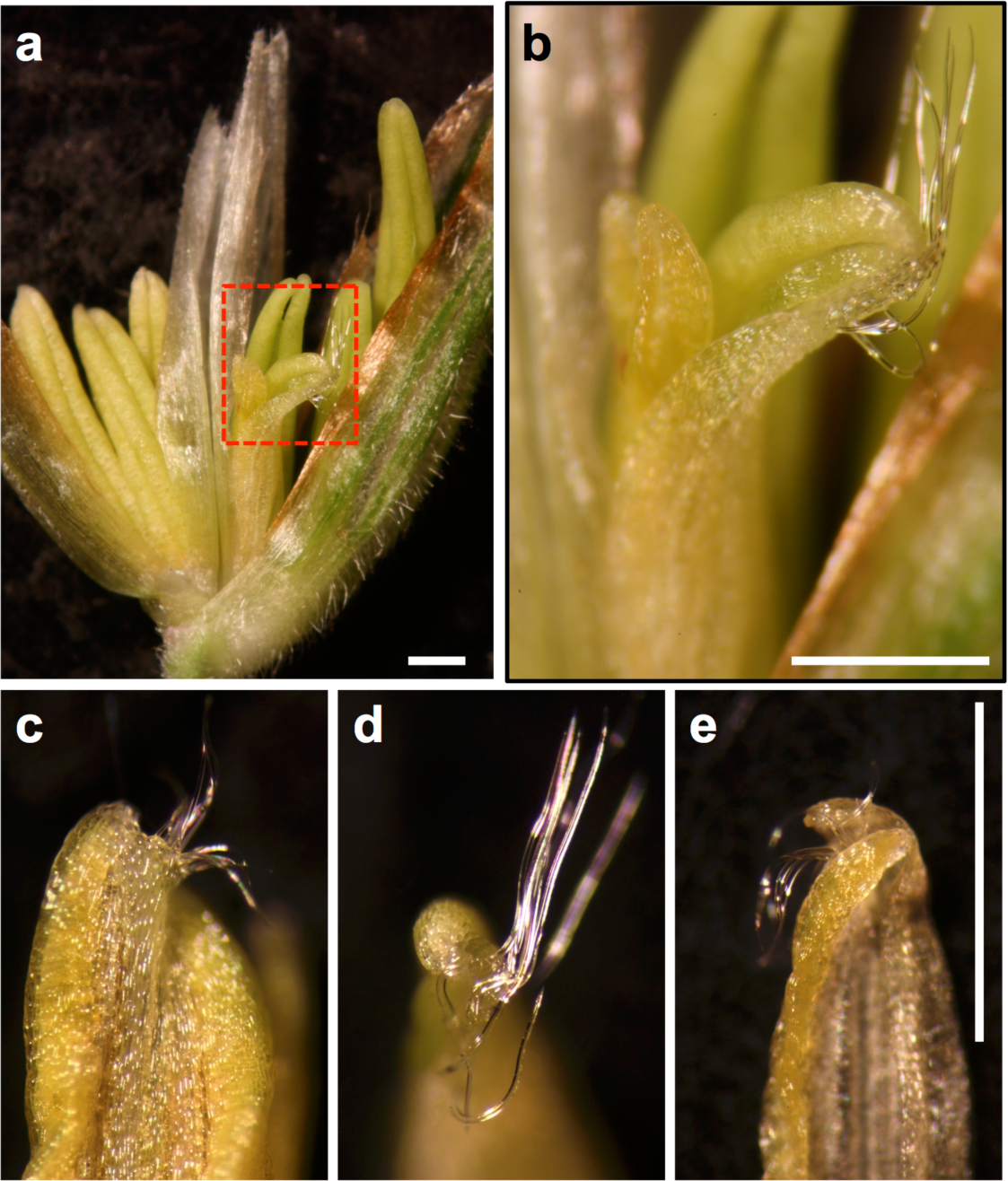
Macrohairs on *drl1; drl2* ectopic anthers. **a,** Dissected staminate floret with ectopic stamen. **b,** Boxed area in (**a**) enlarged to show macrohairs along the apical ridge of the anther. **c-e,** Anthers of ectopic stamens are often amorphic showing deformed theca and empty pollen sacs in addition to macrohairs. Scale bars, 1 mm.

**Figure S6.**
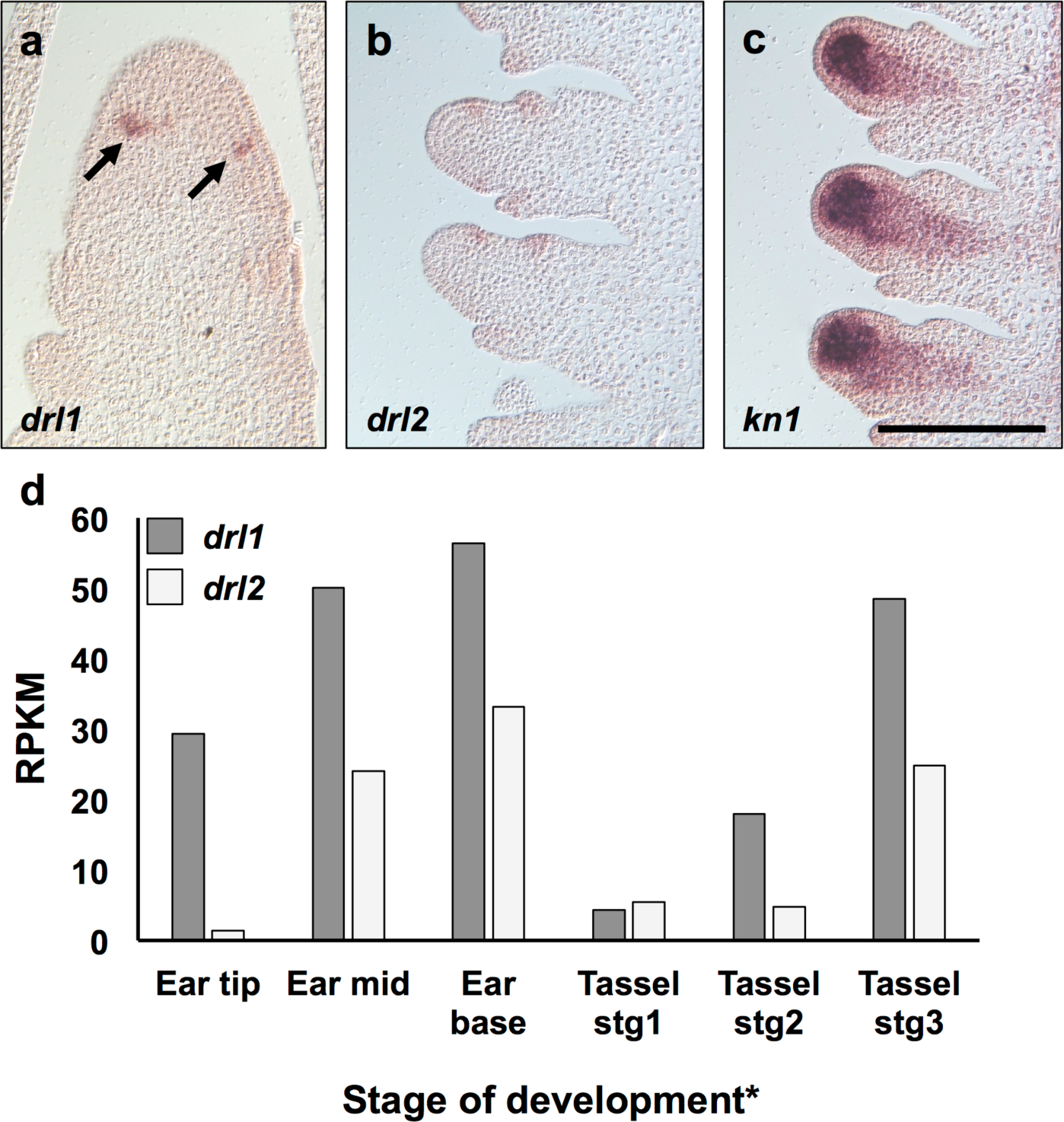
Expression of *drl* genes throughout inflorescence development. **a,** Longitudinal section through IM of a developing ear hybridized with antisense RNA probes to *drl1*; arrow points to *drl1* transcript accumulation in presumptive cryptic bract anlagen. **b,c** Longitudinal sections through early-stage B73 pistillate florets hybridized with antisense RNA probes to *drl2* (**a**) or *kn1* (**b**); Scale bar, 200 µm. **d,** Publicly-available RNAseq data* across B73 developing inflorescences filtered for relative transcript accumulation for *drl1* and *drl2* genes. Ear tip, 1 mm section from the tip of a 10 mm ear (enriched for IM and SPMs); Ear mid, 2 mm section 2 mm from the tip of a 10 mm ear (enriched for SMs); Ear base, 2 mm section 6 mm from the tip of a 10 mm ear (enriched for FMs); Tassel stg1, 1-2 mm; Tassel stg2, 3-4 mm; Tassel stg3, 5-7mm. *www.maizeinflorescence.org.

**Figure S7.**
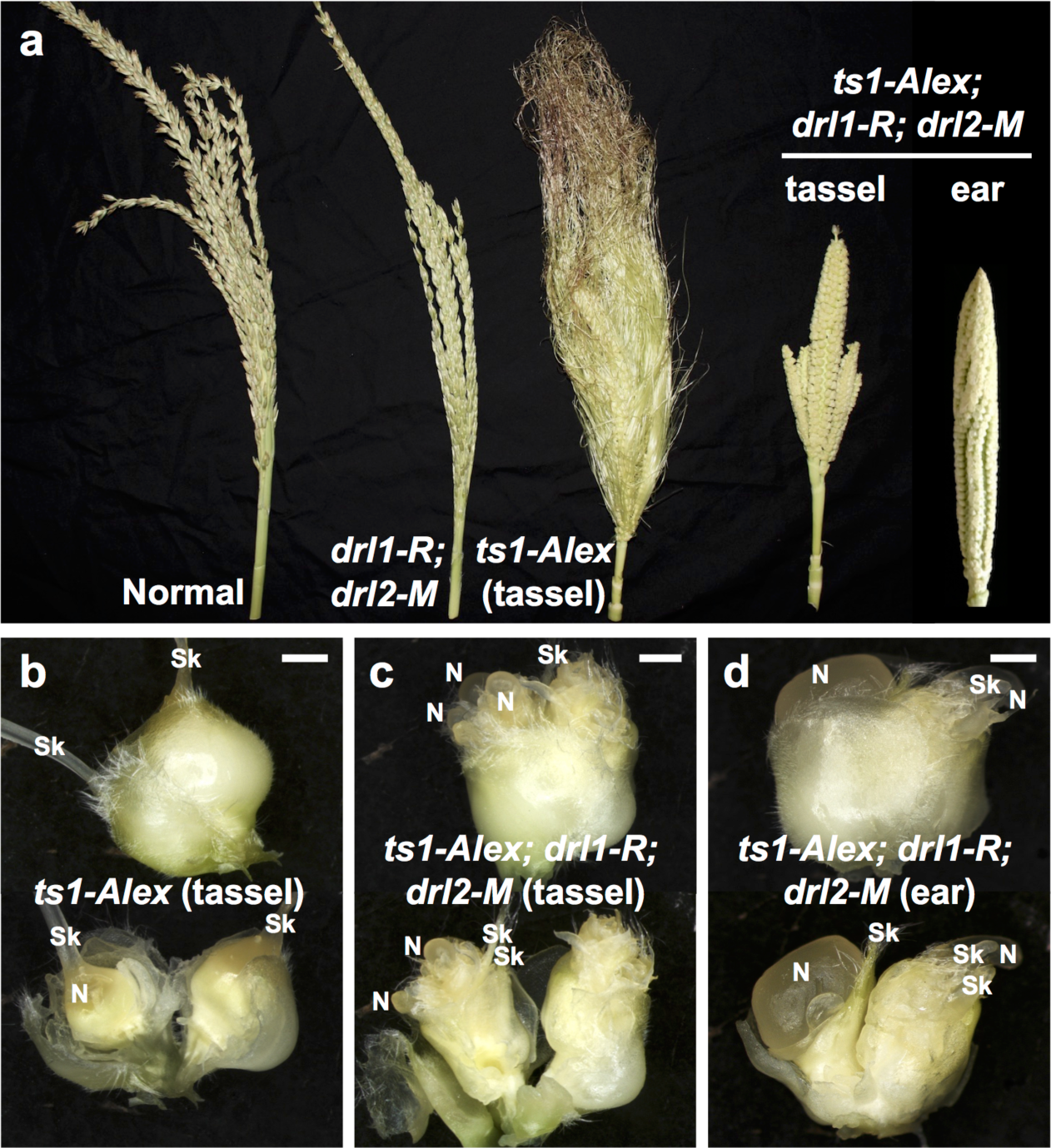
Genetic interaction between *drl1, drl2* and *ts1* mutants. **a,** Mature tassels for normal, *drl1-R; drl2-M*, *ts1-Alex*, and *ts1-Alex; drl1-R; drl2-M* siblings and mature ear for *ts1-Alex; drl1-R; drl2-M* sibling. **b-d,** Dissected florets from *ts1-Alex* (**b**) and *ts1-Alex; drl1-R; drl2-M* (**c**) tassels and from *ts1-Alex; drl1-R; drl2-M* ear (**d**). Glumes were removed in lower panels to expose the inner floret whorls. N, nucellus; Sk, silk. Scale bars, 1 mm.

**Figure S8.**
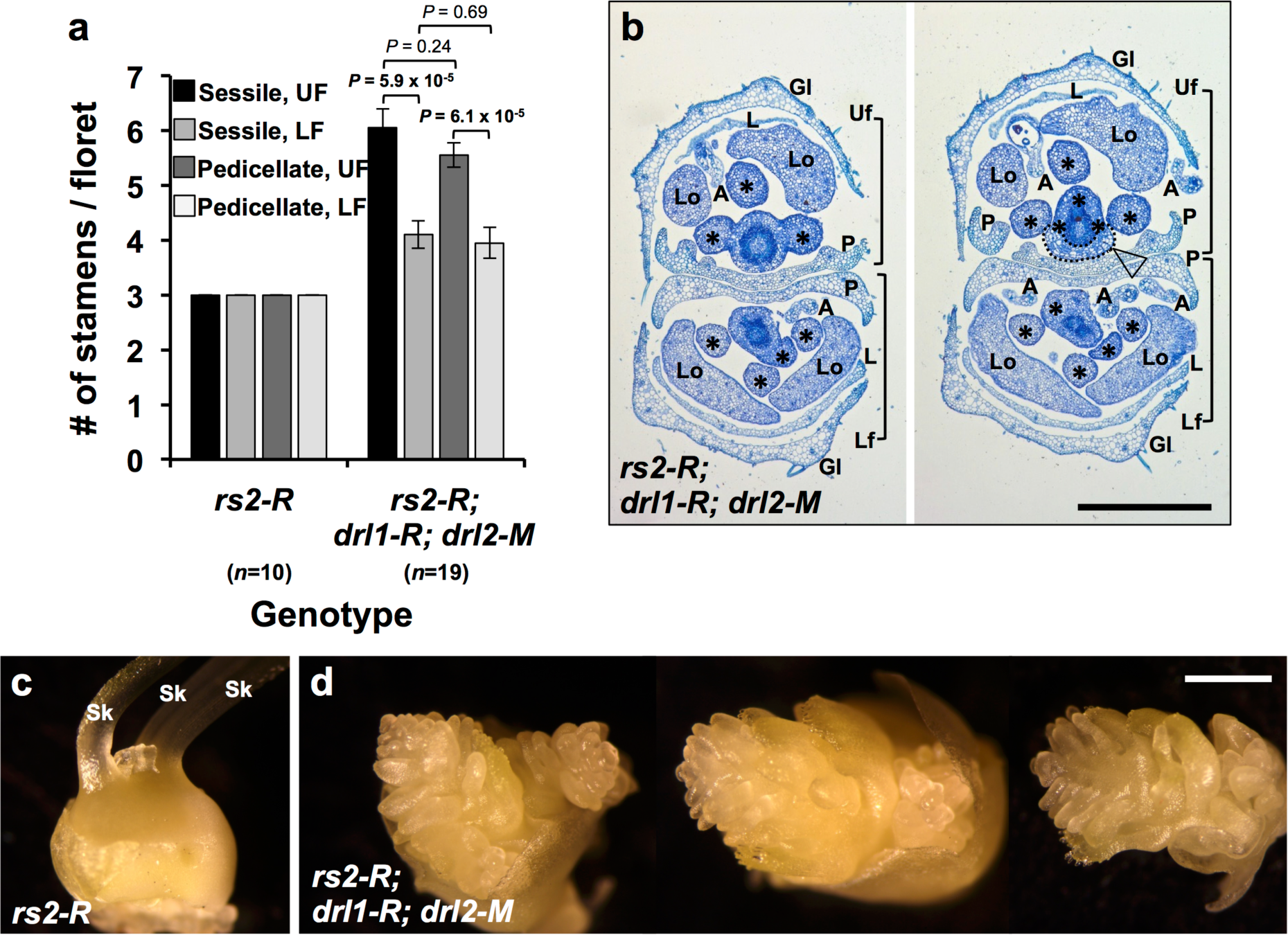
Genetic interaction between *drl1, drl2* and *rs2* mutants. **a,** Quantification of stamens in *rs2-R* and *rs2-R; drl1-R; drl2-M* florets. Mean ±SEM, *P*-values based on two-tailed Student’s *t* tests; *n*, sample size. **b,** Transverse sections of staminate spikelets stained with Toluidine Blue-O taken at base of the mature spikelet (left) and slightly above the base (right). **c,d,** Dissected pistillate spikelet of *rs2-R* (**c**) and *rs2-R; drl1-R; drl2-M* (**d**). Uf, upper floret; Lf, lower floret; L, lemma primordium; P, palea primordium; A, anther primordium; Sk, silk. Asterisks mark stamen primordium. Arrowhead points to ectopic primordium (dotted line) fused to ectopic stamen primordia in the palea axil. Scale bars, 200 µm (**b**) and 1 mm (**c,d**).

